# Structural and Immunological Alterations at the Human Cribriform Plate in *Streptococcus pyogenes* Meningitis: A Case Study

**DOI:** 10.64898/2026.04.28.721468

**Authors:** Jenna Port, Erin Brooks, Jeffery Helgager, Collin Laaker, Melinda Herbath, Matyas Sandor, Zsuzsanna Fabry

## Abstract

*Streptococcus pyogenes* or group A *Streptococcus* (GAS) meningitis is a rare but deadly infection with a high mortality. Its mechanisms of invasion are unknown, but it has been proposed to enter either through the cribriform plate olfactory nerve bundles or the blood brain barrier. Knowledge of how GAS impacts the cribriform plate olfactory nerves can help us better understand GAS pathogenesis and invasion, as well as how it impacts the olfactory nerve bundles. Here we present the case of a 39-year-old otherwise healthy man who presented to the local emergency department with altered mental status and expired the following day. Neuropathologic examination revealed bacterial leptomeningitis; blood and cerebrospinal fluid cultures both grew *Streptococcus pyogenes*. Examination of the cribriform plate was notable for perineural accumulation of GAS around certain olfactory nerve bundles. The accumulation around nerves seems to be random and not correlated to size. Nerves that are impacted by GAS as well as nerves that are not impacted display similar levels of gliosis markers GFAP and podoplanin. Neuropeptide Y, a neuropeptide that implicated in neuro-proliferation and hunger was found to colocalize with CD68 positive immune cells within the nasal epithelium, leading to speculations of its involvement in the inflammatory profile during this case of GAS meningitis. Cribriform plate skull channels had undergone width expansion within the patient, pointing towards local bone marrows involvement during infections. These findings are essential to better understanding the human cribriform plate’s role in CNS immune response and drainage.

## Introduction

*Streptococcus pyogenes*, also referred to as group A *Streptococcus* (GAS) meningitis is a rare infection that accounts for 0.2-1% of all cases of meningitis^1,2^. GAS typically can cause infection in multiple different surfaces of the human body with different severity, ranging from minor diseases such as pharyngitis to severe diseases like sepsis and toxic shock syndrome^3,4^. It has been previously described GAS and other bacteria can potentially invade the CNS either via the cribriform plate or by the blood brain barrier^5,6^. The cribriform plate is a porous structure located in the ethmoid bone in which the olfactory bulb and its nerves. These nerves directly connect the peripheral world to the CNS, leading the cribriform plate to be an important site for immunological regulation, as well as a potential site for bacterial invasion via perineural dissemination. Regardless of bacterial pathways of invasion into the CNS, meningitis can alter olfactory functions, causing anosmia.

Despite knowledge of nerve damage to the olfactory bulbs leading to olfactory dysfunction, there is a lack of understanding of how the olfactory nerves protect themselves from damage due to inflammatory factors from GAS exposure. In this report we present the case of a 39-year-old man with no significant comorbidities who presented to the emergency department with altered mental status. Per family report, he had developed fever and cough over the preceding few days. He ultimately developed radiologic and electroencephalographic evidence of severe anoxic brain injury and expired.

Neuropathologic examination revealed severe global bacterial leptomeningitis involving the cerebral cortex, cerebellum, brainstem, and spinal cord with abundant associated bacteria. Blood and cerebrospinal fluid cultures both grew *Streptococcus pyogenes*. The cause of death was certified as *Streptococcus pyogenes* meningitis with sepsis.

We have found perineural accumulation of GAS around certain olfactory nerves bundles passing through the cribriform plate that does not correlate to circumference. These nerve bundles show increased glial fibrillary acidic protein (GFAP) and podoplanin expression, signifying overall increase inflammation that plagues both nerves with or without GAS accumulation. Finally, in the olfactory nasal epithelium, Neuropeptide Y, an olfactory neuropeptide associated with processing hunger and neuro-proliferation, was colocalized with CD68 positive immune cells.

## Methods

### Specimen retrieval & Processing

Upon harvesting, the cribriform plate was placed into 4% formalin for 24 hours. After 24 hours, the cribriform plate was placed into a decalcification solution to decalcify any bone prior to sectioning for 2 weeks, swapping out the solution every other day. Post decalcification, the sample was placed into a sucrose solution for 24 hours. The sample was then frozen in optimal cutting temperature (OCT) compound. The sample was then serial sectioned in a cryostat coronally at 40 microns.

### Immunohistochemistry

Sections were blocked with a 3% BSA 5% Normal Donkey Serum with 0.3% Tritonx100 solution for 1 hour. Primary antibodies of Streptococcus group A, Neuropeptide Y, CD45, CD3, CD68, CD19, GFAP, and Podoplanin were incubated overnight at 4 degrees Celsius. Slides were washed with a BSA diluted in PBS, then stained with secondary antibodies Streptavin 488, Donkey anti rabbit 568, Donkey anti mouse 647, Donkey anti chicken 647, or donkey anti rat 647 for 1 hour at room temperature. Slides were then washed, dried, and slide covers were mounted with hardest mounting media with DAPI. Imaging occurred on a confocal laser scanning microscope Olympus FV1-ASW Fluoview.

## Results

### Accumulation of bacteria around nerves is specific to streptococcus group A meningitis patient

IF-IHC staining of TUBB3 and Streptococcus group A showed significant accumulation of strep pyogenes around TUBB3+ nerve bundles (Figure 1A). The accumulation of bacteria occurs is not dependent on the size of the nerve bundle as size did not significantly correlate to perineural strep pyogenes accumulation (Figure 1). It has been theorized that the pathogenesis of bacterial meningitis can be traced to entering the central nervous system via the nasal lymphatic system through the cribriform plate. Although this has never been confirmed in cases of streptococcus pyogenes. Secondly, the cribriform plate is known as a site of CSF drainage that can fluctuate depending on the disease/ inflammatory context^7–9^. Previous studies in mice have shown the potential of bacteria to hijack the nasal lymphatics as an access point to the CNS, and have traveled through the cribriform plate perineurally as a route to the CNS^10^. of strep pyogenes perineurally proves that there is potential involvement, either allowing strep pyogenes to infiltrate the CNS or drain from the CNS utilizing the cribriform plate and the olfactory nerves passing through it. Further studies to examine the pathogenesis and migration of streptococcus pyogenes to or from the CNS will need to be conducted to give a definitive answer to these speculations.

**Figure 1.**
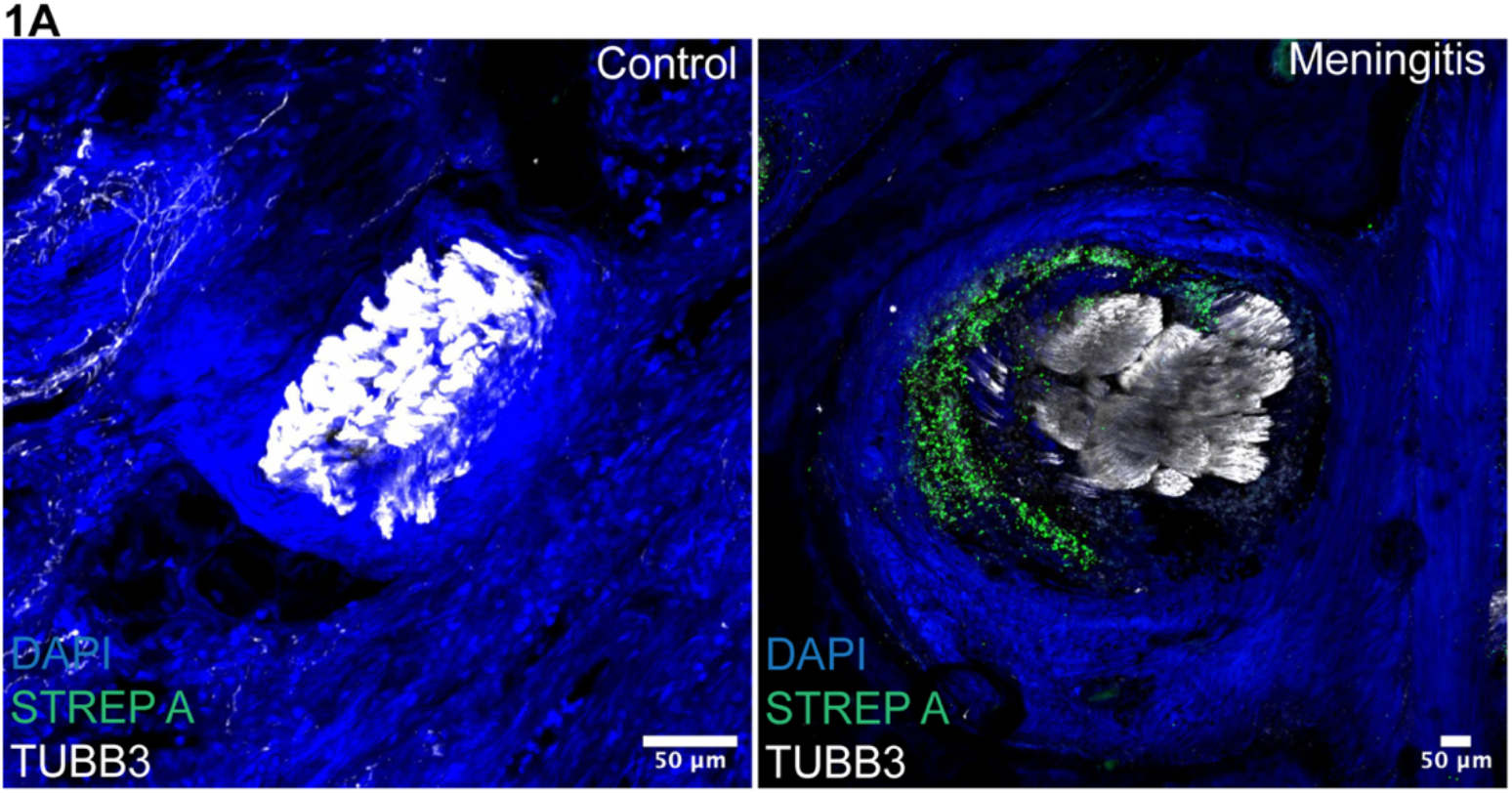
Group A streptococcus accumulates perineurally around olfactory nerves at the human cribriform plate. **1A)** representative images of human olfactory nerves in a control specimen versus the GAS meningitis patient. Stained with Dapi (blue), Group A Strep (green), Tubb3 (white).

### Olfactory nerve bundles express similar amounts of gliosis markers despite being exposed to streptococcus pyogenes or not

Based off the previous finding of perineural accumulation of group A streptococcus, we questioned why streptococcus pyogenes was only accumulating on certain olfactory nerve bundles, rather than others. We hypothesized that there could be potential protective measures that the olfactory nerve bundles. To test this hypothesis, we stained for gliosis markers, as well as neuropeptides. There are significant increases in podoplanin and GFAP within the nerves of the meningitis patient, signifying increased gliosis and inflammation (Figure 2A-D). Within glial cells, it is known that they express neuropeptides dependent on the region in the brain that the cells are located in^13^. Further, NPY is shown to be involved in neuro-proliferative measures that can be associated with repair to olfactory ensheathing cells and spinal cords^14^. To examine the neuropeptide presence within the olfactory nerve bundles we stained for neuropeptide Y, a peptide that is highly involved in olfactory functions. There was no significant difference in NPY expression in nerves impacted by streptococcus pyogenes versus ones that are not impacted by bacteria.

**Figure 2.**
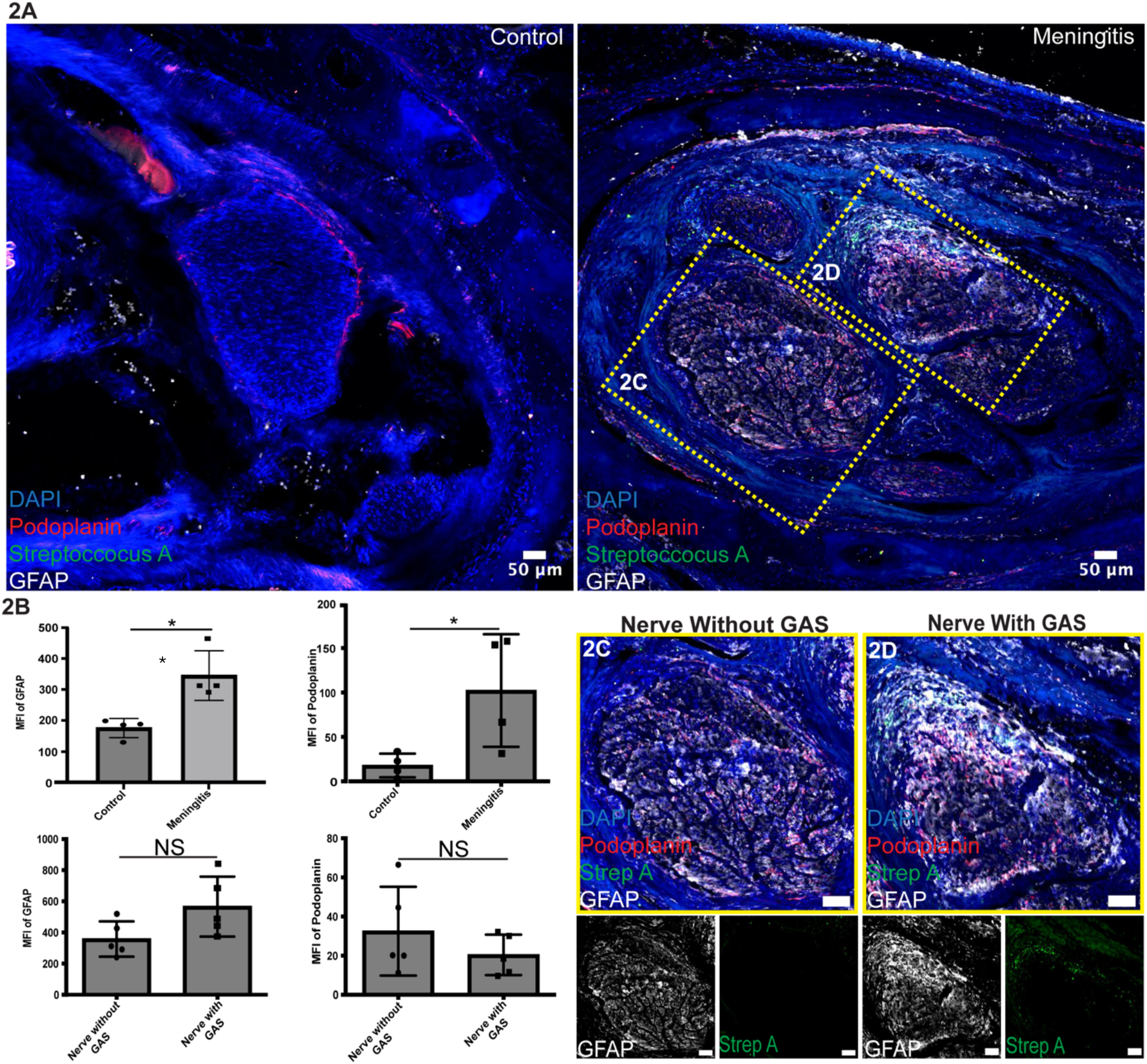
Significant upregulation of gliosis markers in GAS meningitis case occurs, but nerves impacted by GAS do not have increased gliosis in comparison to nerves not impacted by GAS. **2A** Representation of gliosis markers GFAP and podoplanin increasing in the olfactory nerves of the GAS patient in comparison to a non-CNS impacted control. **2B** Quantitative analysis of mean fluorescent intensity (MFI) of gliosis markers. Unpaired T-test shows (Top right) GFAP MFI control vs. meningitis P= 0.0075, (Top left) Podoplanin MFI control vs. meningitis P=0.0402, (Bottom left) GFAP MFI nerve without GAS vs. nerve with GAS P=0.0705, (Bottom right) Podoplanin MFI nerve without GAS vs. nerve with GAS P=0.3093. Significance portrayed as: n.s. = nonsignificant, p > 0.05; *p < 0.05; **p < 0.01; ***p < 0.001; ****p < 0.0001. **2C** Graphic representation of a nerve from the GAS meningitis patient without GAS accumulation and its relative GFAP and Podoplanin expression. **2D** Graphic representation of a nerve from the GAS meningitis patient with GAS accumulation and its relative GFAP and Podoplanin expression.

### Colocalization of neuropeptide Y with immune cells within the cribriform plate nasal epithelium during meningitis

Incidentally, upon examination of the olfactory nasal epithelium we have found that there is significant colocalization of MFI between CD45+ positive cells and NPY. With this colocalization of immune cells, this could implicate potential interactions between NPY and immune cells (Figure 3A). To gain more insight on specific immune cell subtypes interacting with NPY, we conducted IF-IHC for T cells (CD3) (Figure 3C), B cells (CD19) (Figure 3D), and myeloid cells (CD68) (Figure 3B). We found that CD68+ cells colocalize most closely with NPY in comparison to other cell types. Previously, NPY’s involvement with immune cells has been confirmed in cases of multiple sclerosis.

**Figure 3.**
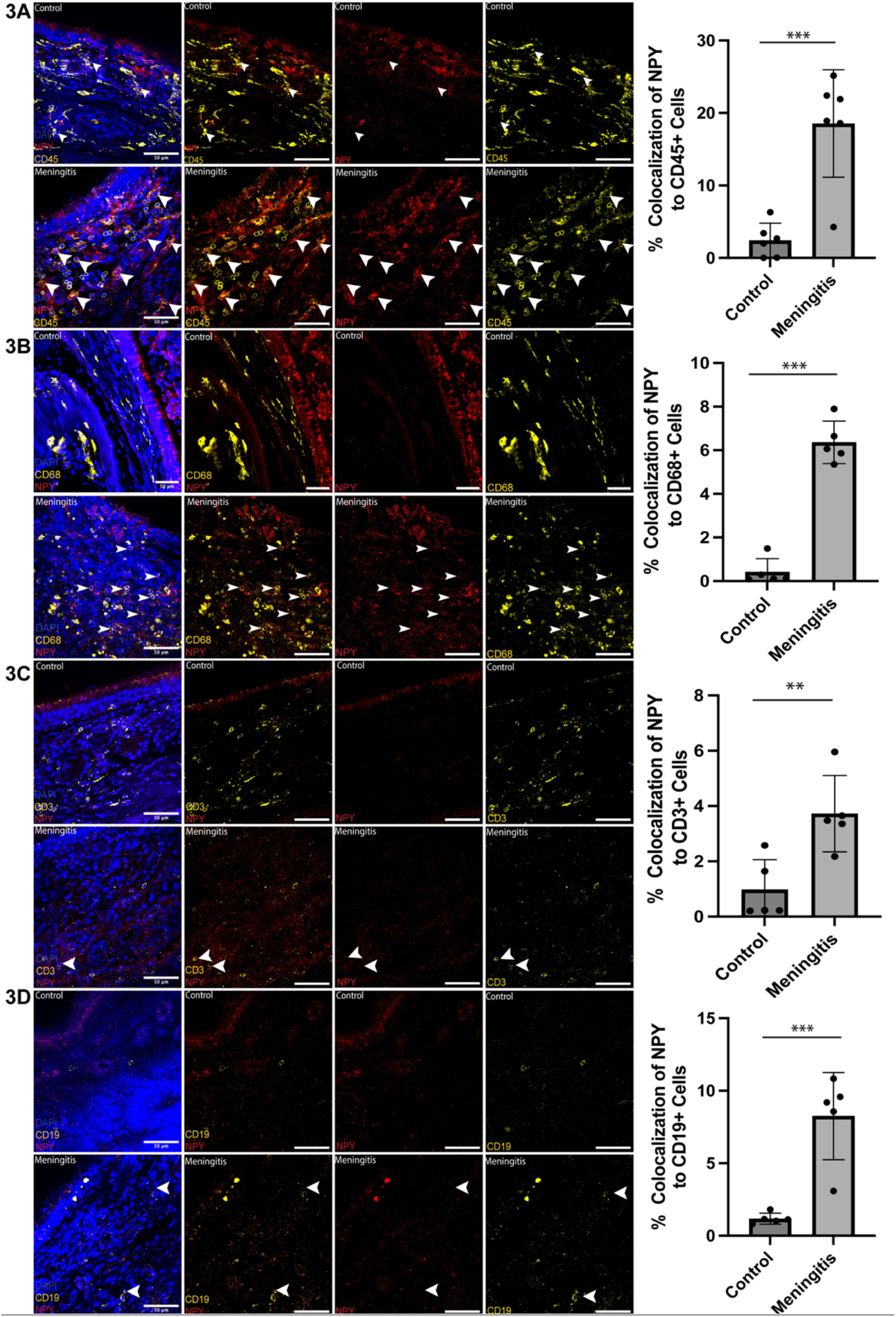
Colocalization of neuropeptide Y with immune cells within the cribriform plate nasal epithelium during meningitis. **3A** Representative image of CD45+ leukocytes colocalizing with neuropeptide Y (NPY) in meningitis patient nasal epithelium along with quantification of % colocalization of CD45+ cells to NPY, unpaired T Test comparing control vs. meningitis P=0.0005. **3B** representative images of CD68+ myeloid cells colocalizing with neuropeptide Y (NPY) in meningitis patient nasal epithelium along with quantification of % colocalization of CD68+ cells to NPY, unpaired T Test comparing control vs. meningitis P<0.0001. **3C** representative images of CD3+ T cells colocalizing with neuropeptide Y (NPY) in meningitis patient nasal epithelium along with quantification of % colocalization of CD3+ cells to NPY, unpaired T Test comparing control vs. meningitis P=0.0081. **3D** representative images of CD19+ B cells colocalizing with neuropeptide Y (NPY) in meningitis patient nasal epithelium along with quantification of % colocalization of CD19+ cells to NPY, unpaired T Test comparing control vs. meningitis P=0.0008. Significance portrayed as: n.s. = nonsignificant, p > 0.05; *p < 0.05; **p < 0.01; ***p < 0.001; ****p < 0.0001.

### Expansion of skull channel vessels within the cribriform plate during GAS infection

Previously we have determined that skull channels are present within the murine cribriform plate and have access to cerebrospinal fluid sampling from perineural areas. Further, we found that mycobacterium tuberculosis can disseminate into skull bone marrow channels and further, the bone marrow with it, proving the intricate capabilities of bacterial invasion. To expand on this finding, we observed bone marrow channels within the GAS meningitis cribriform plate. Immunohistochemical staining of CD31, streptococcus pyogenes, and DAPI shows presence of GAS bacteria within the skull channel and coagulated within the bone marrow (FIGURE 4A). Further, there are significant increases of the width of skull bone marrow channel vessels, along with the bone near it (unpaired t-test p<0.0315) (FIGURE, 4B-C). This is the first documented response of expansion of skull bone marrow channels due to meningitis infection as well as skull bone marrow channels The skull bone marrow is known to be a dynamic expanding niche ^11^It has been previously reported that skull bone marrow and vessels within the calvaria do expand with age, potentially due to pro-inflammatory cytokines ^12^. These findings contribute to the narrative that bone marrow can alter its methods of expansion in response to bacteria such as GAS.

**Figure 4.**
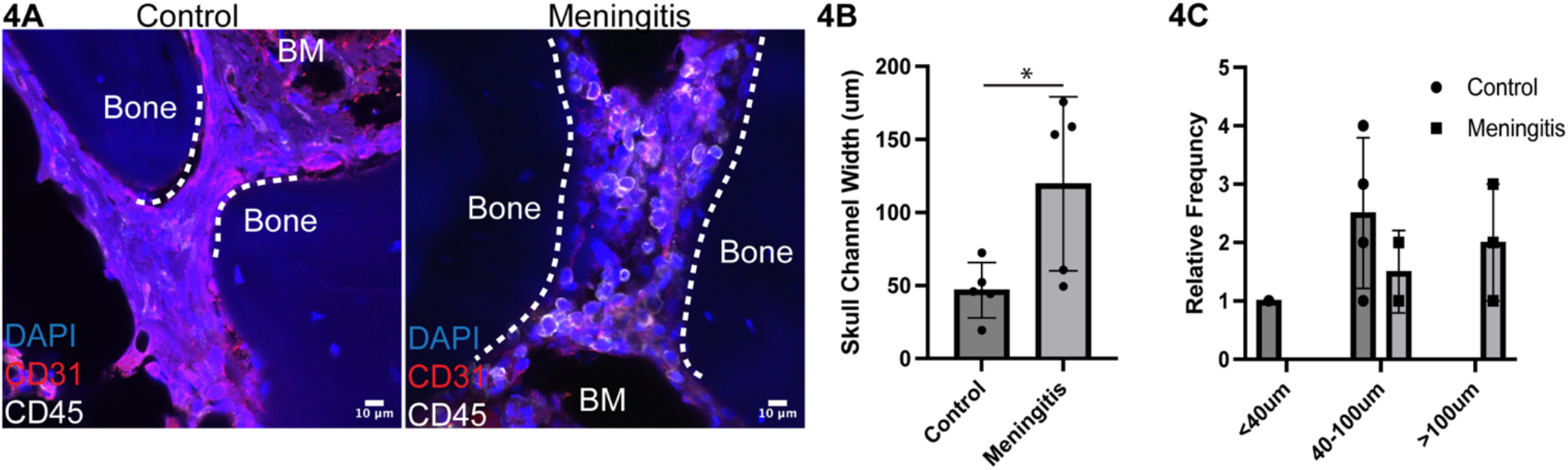
Expansion of skull channel vessels within the cribriform plate during GAS infection. **4A** Graphic representation of a non-CNS disease control cribriform plate skull channel and the GAS meningitis patient cribriform plate skull channel. **3B** Quantitative analysis measuring the skull width of control vs. GAS meningitis patient. Unpaired T-test P=0.0315. Significance portrayed as: n.s. = nonsignificant, p > 0.05; *p < 0.05; **p < 0.01; ***p < 0.001; ****p < 0.0001. **3C** Relative frequency of skull bone channels of <40 um width, 40-100 um width, and >100 um width in control and meningitis patients.

## Discussion

GAS meningitis is a bimodal disease particularly impacting the young and the old, however middle aged individuals are still at risk^3^. Investigating the impact of meningitis at the cribriform plate will prove its function in either drainage of the infected CSF, or perhaps bacteria infiltration through the nasal lymphatics and perineural routes. Another feature that will need to be studied in future cases of cribriform plate is the appearance of randomized accumulation of GAS perineurally in the olfactory nerve bundles. Based off of our serial sections, there seems to be no logical explanation as to why certain olfactory nerves have accumulations of GAS, and there are no structural differences between non impacted versus impacted nerves. Studying more cases of GAS meningitis at the cribriform plate would allow us to better gage whether this accumulation is random, or if the GAS targets certain olfactory nerve bundles versus others. This confirms that perineural accumulation of GAS and perhaps other bacteria does occur in humans, but further studies are needed to confirm how the olfactory nerves damage from GAS alters CNS outflow, and olfactory function.

Neuropeptides have been known to partake in host functions during meningitis^15–19^. In other forms of bacterial meningitis have been found to manipulate other nociceptors such as calcitonin gene receptor peptide (CGRP) and substance P in weakening immune responses and the overall host defense in other forms of bacterial meningitis^17,20^. Further, NPY has been found to increase in the brain during pneumococcal meningitis, however NPY’s function in bacterial meningitis, and more specifically, GAS meningitis remains unknown. Our finding of NPYs colocalization to CD68 positive immune cells may implicate its ability to modulate immune cells during GAS meningitis. NPY has been implicated in modulating messenger immune cells, primarily macrophages and dendritic cells to dependent on disease context^21–25^. We overall are limited in our methodology that we could perform on this postmortem tissue. Although IF-IHC remains a reliable method for visualizing the structural features and immune cell presence of the cribriform plate, expanding to other forms of experimentation such as single cell RNA sequencing, flow cytometry, and radioimmunoassay’s would significantly help prove true interactions between CD68 positive cells and NPY, and what this does to alter immune cell phenotypes and function in the context of GAS meningitis.

## Conclusions

We have found the novel finding of interactions with NPY and CD68 positive immune cells at the cribriform plate, furthermore in-depth studies need to be conducted to confirm these interactions and to study the mechanisms as to why this occurs. GAS has capability to disseminate into the skull bone marrow, and skull bone marrow channels width expands with GAS meningitis. Structural differences in the olfactory nerve bundles are not influenced by the presence of GAS, and GAS’s perineural accumulation leads to novel proof that the perineural pathway is utilized within human cribriform plates.

## Bibliography

1. Schlech WF Iii, Ward JI, Band JD, Hightower A, Fraser DW, Broome CV. Bacterial Meningitis in the United States, 1978 Through 1981: The National Bacterial Meningitis Surveillance Study. JAMA. 1985;253(12):1749–1754. doi:10.1001/jama.1985.03350360075022

2. van de Beek D, de Gans J, Spanjaard L, Sela S, Vermeulen M, Dankert J. Group A Streptococcal Meningitis in Adults: Report of 41 Cases and a Review of the Literature. Clin Infect Dis. 2002;34(9):e32–e36. doi:10.1086/339941

3. Thacharodi A, Hassan S, Vithlani A, et al. The burden of group A Streptococcus (GAS) infections: The challenge continues in the twenty-first century. iScience. 2025;28(1). doi:10.1016/j.isci.2024.111677

4. Newberger R, Hollingshead CM. Group A Streptococcal Infections. In: StatPearls. StatPearls Publishing; 2025. Accessed March 20, 2025. http://www.ncbi.nlm.nih.gov/books/NBK559240/

5. Cutforth T, DeMille MM, Agalliu I, Agalliu D. CNS autoimmune disease after Streptococcus pyogenes infections: animal models, cellular mechanisms and genetic factors. Future Neurol. 2016;11(1):63. doi:10.2217/fnl.16.4

6. Dileepan T, Smith ED, Knowland D, et al. Group A Streptococcus intranasal infection promotes CNS infiltration by streptococcal-specific Th17 cells. J Clin Invest. 2015;126(1):303–317. doi:10.1172/JCI80792

7. Hsu M, Rayasam A, Kijak JA, et al. Neuroinflammation-induced lymphangiogenesis near the cribriform plate contributes to drainage of CNS-derived antigens and immune cells. Nat Commun. 2019;10(1):229. doi:10.1038/s41467-018-08163-0

8. Hsu M, Laaker C, Madrid A, et al. Neuroinflammation creates an immune regulatory niche at the meningeal lymphatic vasculature near the cribriform plate. Nat Immunol. 2022;23(4):581–593. doi:10.1038/s41590-022-01158-6

9. Choi YH, Hsu M, Laaker C, et al. Dual role of Vascular Endothelial Growth Factor-C (VEGF-C) in post-stroke recovery. Published online September 1, 2023:2023.08.30.555144. doi:10.1101/2023.08.30.555144

10. Dileepan T, Smith ED, Knowland D, et al. Group A Streptococcus intranasal infection promotes CNS infiltration by streptococcal-specific Th17 cells. J Clin Invest. 2016;126(1):303–317. doi:10.1172/JCI80792

11. Liu L, Zhang X, Chai Y, Zhang J, Deng Q, Chen X. Skull bone marrow and skull meninges channels: redefining the landscape of central nervous system immune surveillance. Cell Death Dis. 2025;16(1):1–17. doi:10.1038/s41419-025-07336-2

12. Koh BI, Mohanakrishnan V, Jeong HW, et al. Adult skull bone marrow is an expanding and resilient haematopoietic reservoir. Nature. 2024;636(8041):172–181. doi:10.1038/s41586-024-08163-9

13. Harvey JD, Heinbockel T. Neuromodulation of Synaptic Transmission in the Main Olfactory Bulb. Int J Environ Res Public Health. 2018;15(10):2194. doi:10.3390/ijerph15102194

14. Hansel DE, Eipper BA, Ronnett GV. Neuropeptide Y functions as a neuroproliferative factor. Nature. 2001;410(6831):940–944. doi:10.1038/35073601

15. Täuber MG, Ferriero D, Kennedy SL, Sheldon RA, Guerra-Romero L. Brain Levels of neuropeptide Y in experimental pneumococcal meningitis. Mol Chem Neuropathol. 1993;18(1):15–26. doi:10.1007/BF03160019

16. Lu YZ, Nayer B, Singh SK, et al. CGRP sensory neurons promote tissue healing via neutrophils and macrophages. Nature. 2024;628(8008):604–611. doi:10.1038/s41586-024-07237-y

17. Li Y, Wang L, Gao Z, et al. Neuropeptide Calcitonin Gene-Related Peptide Promotes Immune Homeostasis of Bacterial Meningitis by Inducing Major Histocompatibility Complex Class II Ubiquitination. J Infect Dis. 2024;229(3):855–865. doi:10.1093/infdis/jiad358

18. Kumar P, Williams JN, Durkin KL, et al. Neuropeptide α-MSH exerts pro-inflammatory eCects on Neisseria meningitidis infection in vitro. Inflamm Res. 2010;59(2):105–113. doi:10.1007/s00011-009-0076-9

19. Gao F, Hu H. Nociceptors and Macrophages in Bacterial Meningitis: Partners in Crime? Neurosci Bull. 2024;40(3):418–420. doi:10.1007/s12264-023-01141-7

20. Liu H, Kong X, Zeng Y, et al. From pain to meningitis: bacteria hijack nociceptors to promote meningitis. Front Immunol. 2025;15. doi:10.3389/fimmu.2024.1515177

21. Zhu P, Sun W, Zhang C, Song Z, Lin S. The role of neuropeptide Y in the pathophysiology of atherosclerotic cardiovascular disease. Int J Cardiol. 2016;220:235–241. doi:10.1016/j.ijcard.2016.06.138

22. Li C, Wu X, Liu S, Zhao Y, Zhu J, Liu K. Roles of Neuropeptide Y in Neurodegenerative and Neuroimmune Diseases. Front Neurosci. 2019;13. doi:10.3389/fnins.2019.00869

23. Ferreira-Marques M, Carmo-Silva S, Pereira J, et al. Restoring neuropetide Y levels in the hypothalamus ameliorates premature aging phenotype in mice. GeroScience. Published online February 27, 2025. doi:10.1007/s11357-025-01574-0

24. Buttari B, Profumo E, Domenici G, et al. Neuropeptide Y induces potent migration of human immature dendritic cells and promotes a Th2 polarization. FASEB J. 2014;28(7):3038–3049. doi:10.1096/fj.13-243485

25. Yi M, Li H, Wu Z, et al. A Promising Therapeutic Target for Metabolic Diseases: Neuropeptide Y Receptors in Humans. Cell Physiol Biochem. 2017;45(1):88–107. doi:10.1159/000486225

